# Species-Agnostic Transfer Learning for Cross-species Transcriptomics Data Integration without Gene Orthology

**DOI:** 10.1101/2023.08.11.552752

**Authors:** Youngjun Park, Nils Paul Muttray, Anne-Christin Hauschild

## Abstract

Novel hypotheses in biomedical research are often developed or validated in model organisms such as mice and zebrafish and thus play a crucial role, particularly in studying disease mechanisms and treatment responses. However, due to biological differences between species, translating these findings into human applications remains challenging. Moreover, commonly used orthologous gene information is often incomplete, particularly for non-model organisms, and entails a significant information loss during gene-id conversion. To address these issues, we present a novel methodology for species-agnostic transfer learning with heterogeneous domain adaptation. We built on the cross-domain structure-preserving projection and extended the algorithm toward out-of-sample prediction, a common challenge in biomedical sequencing data. Our approach not only allows knowledge integration and translation across various species without relying on gene orthology but also identifies similar GO biological processes amongst the most influential genes composing the latent space for species integration. Subsequently, this enables the identification and functional annotation of genes missing from public orthology databases. Finally, we evaluated our approach with four different single-cell sequencing datasets focusing on out-of-sample prediction and compared it against related machine-learning approaches. In summary, the developed model outperforms all related methods working without prior knowledge when predicting unseen cell types based on other species’ data. The results demonstrate that our novel approach allows knowledge transfer beyond species barriers without the dependency on known gene orthology but utilizing the entire gene sets.

## 1 Introduction

Model organisms such as mice and zebrafish play a crucial role in developing and validating new hypotheses in biomedical research, particularly in studying disease mechanisms and treatment responses. However, due to biological differences between species translating these findings into human applications remains challenging. Therefore, additional intensive experimental validation is required to confirm whether the knowledge gained from one species can be translated into humans[1]. Although researchers may generate similar data on different model organisms during this process, it is often infeasible to integrate and analyze due to data heterogeneity. This data heterogeneity originates from different sources, such as signal noise, batch effects, technological heterogeneity, and biological heterogeneity. This presents a challenging task in data integration, requiring either domain-specific noise handling, a multi-modal analysis methodology capable of handling different data types [2], or a cross-species transfer learning methodology [3]. In biomedical research, biological heterogeneity where each species has a different set of genes responsible for certain biological functions is a critical hurdle to be addressed. The current approach to handle this issue is to utilize gene orthology information. In this inter-species data integration, biological heterogeneity necessitates a gene-id conversion with human-curated data such as an orthologous gene. However, this approach faces two major challenges: a many-to-many mapping for various gene families and severe information loss during the gene-id conversion.

Cell-type label projection is the main task for the recent transfer learning model with single-cell RNA sequencing data. The recent bone-marrow cell type classification study with mice and humans found that transfer learning can improve classification accuracy when the target data is limited in a small sample size [4]. They utilized orthologous gene information, representing meta-information about genes in two species. scAdapt [5] implemented an adversarial domain adaptation network for the cross-species label projection. CAME [6] applied a graph neural network to exploit orthology information fully. However, they still rely on the orthologous gene information. In the scETM (Single-Cell Embedded Topic Model) work, the features of two different species are transformed by orthologous gene pairs. Although this work showed the possibility of cross-species transfer learning, this is limited to homogeneous domain transfer learning using orthologous gene set information [7]. SAMap aims to integrate various transcriptomes from different species by exploiting gene orthology information. With the graph-based method, they infer cell types across different species at the Metazoa level. This algorithm transfers labels through the species-barrier and considers evolutionary trajectory by considering many-to-many orthology relations between species [8]. Moreover, it enables the analysis of cell-type families in different species. This work has improved the gene-id conversion method to have less information loss. However, it still relies on gene orthology information. scGEN employed *Variational Autoencoder* (VAE) and feature-vector arithmetics. It was able to predict similar cell perturbation responses across different species [9]. However, they also used orthologous gene information to match different genes on different species for training the model. Transfer learning approaches are commonly differentiated into two categories: (1) homogeneous and (2) heterogeneous domain adaptation. In homogeneous transfer learning the domains share the same feature space but have different distributions. This is the case, when gene ID can be mapped to the same ids in a different species with gene orthology, for instance. Heterogeneous transfer learning, in contrast, supposes that the domains have different feature spaces and distributions. Thus, it can allow for knowledge integration and translation across various species’ datasets without gene-ID conversion relying on external knowledge.

To facilitate this, we developed a novel approach called *Species-Agnostic Transfer Learning* (SATL) that identifies a common latent space of different species without relying on external knowledge. The approach builds on the *Cross-Domain Structural Preserving Projection* (CDSPP) method where the model learns a projection matrix for a domain-invariant feature subspace to reduce the discrepancy between domains [10]. It allows to incorporate the entire dataset in the cross-species analysis.

## 2 Results

While there is a large set of bioinformatics and data science methodologies that allow the data-driven integration of biological and biomedical datasets, data-driven knowledge transfer between species only recently became possible with novel transfer learning approaches. However, most of these studies have a limitation: they need to employ external data to homogenize heterogeneous feature spaces in order to transfer learned models amongst different domains. One of the biggest issues with this methodology is the inevitable loss of information during the feature conversion based on gene orthology information [11–14]. Thus we are presenting a new methodology to investigate biological knowledge in a completely data-driven way without severe information loss.

### 2.1 SATL adapts heterogeneous domains of feature spaces from different species

The main goal of our work is to explore an alternative approach to transfer classifiers beyond the barriers between species. Therefore, we investigated different *Heterogeneous Domain Adaptation* (HDA) methodologies and related fields such as *Domain Adaptation* (DA) and *Generalized Zero-Shot Learning* (GZSL) for their capabilities to model shared cell types in single-cell sequencing data across two species. In contrast to the current approaches, no information about orthologous gene will be used. Instead, the newly developed SATL method will align the data space of the source and target domain with the aid of a subset of labeled data from the target domain and source domain. A detailed description of the methodology is provided in the supplementary material.

The main idea of SATL is to extend a semi-supervised HDA algorithm to enable GZSL on heterogeneous biomedical data sets, such as single-cell sequencing data from different species, for instance. The evaluation scheme, therefore, follows the strategy of a transductive GZSL setting, where the labels of a certain fraction of classes of the target species are unknown during training. In the following results, we demonstrate and evaluate cross-species transfer learning. At first, with lps-stimulated macrophage dataset from the four species, we show how the proposed method works and how could the result be interpreted in a biologically meaningful way by functional analysis. SATL is compared with *Mutual Nearest Neighbors* (MNN), 2.2, and identified latent space is investigated, 2.3. After that, we further evaluated the performance of SATL with other models having a similar concept and purpose. Three human-mouse pair datasets are used for this comparison. This comparison works contain comparisons to GZSL models and DA 2.4 methods. The detailed results from three single-cell sequencing data are also investigated 2.5.

### 2.2 SATL outperformed mutual nearest neighbors in heterogeneous domain adaptation task

Here, we explore the performance of SATL in comparison to MNN, a popular method for single-cell data integration and batch effect mitigation. While MNN is intended for homogeneous domain adaptation, requiring the same order and genes, the methodology is closely related to CDSPP. For the cross-species integrative analysis without gene-ID conversion, we first extract features from the species with different numbers of genes. These features consist of different latent spaces for each species dataset, each corresponding to heterogeneous domains. We compared SATL to MNN in a heterogeneous domain adaptation task, with the lps-stimulated macrophage dataset [15].

Previously, scGEN has shown that vector arithmetics on latent space can predict gene signatures of unseen cell types in different species in the lps-stimulated macrophage dataset [9]. Likewise, we analyzed SATL on the same dataset. The stimulation dataset has labels about experimental conditions instead of cell types. Training and testing are done in the same procedure as other single-cell datasets. There are four species and each species has four labels, unst, lps2, lps4, and lps6. Compared to the cross-species analysis of the original work, where only 2,336 genes were investigated [16], and the scGEN study, where 6,619 genes were investigated [9], our analysis can target all expressed genes from both species, see Methods for details. In this analysis, analyzed genes were 10,000 to 13,000 genes after single-cell RNA data preprocessing (Table 4).

Here, one group of cells from target species is masked and predicted based on the model trained with data from source species. The evaluation is done in all pairwise scenarios among all four species. At first, we extracted features using *Principal Component Analysis* (PCA) and applied MNN correction to mitigate batch effects and align the feature spaces, assuming that the two species had homogeneous domains. Subsequently, we trained a *Random Forest* (RF) model on the processed source species data to predict cell types of target species (Figure 2). Simultaneously, SATL was employed on the same features extracted from PCA. In most of the scenarios, our SATL model predicts the masked cells with the correct label. However, the comparison between both approaches shows that, the RF model trained with MNN-corrected features, designed for homogeneous domain adaptation, cannot correct heterogeneous domains in cross-species data integration tasks. In most of the scenarios, it failed to predict the masked cell type of the target species. Additionally, we extracted *Locality Preserving Projection* (LPP) features instead of PCA features and combined them with the MNN and RF classification approach. However, no improvement in prediction was achieved.

**Fig. 1.**
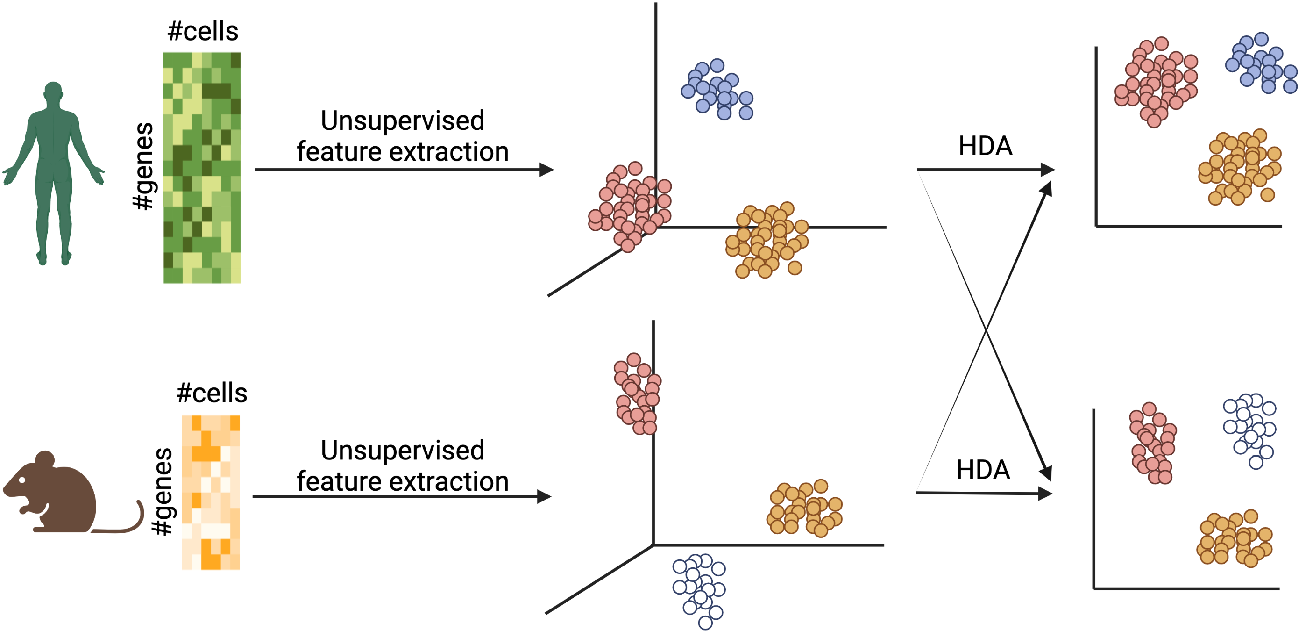
Overview of cross-species single-cell data integration and label-projection utilizing the developed SATL workflow. Different species’ gene expression profiles can be used without gene-id conversion. Extracted features from each species are not in the same domain space. Heterogeneous domain adaptation (HDA) aligns these different domains into a common latent space. In this common space, unlabelled data in the target species can be labeled with knowledge of the source species.

**Fig. 2.**
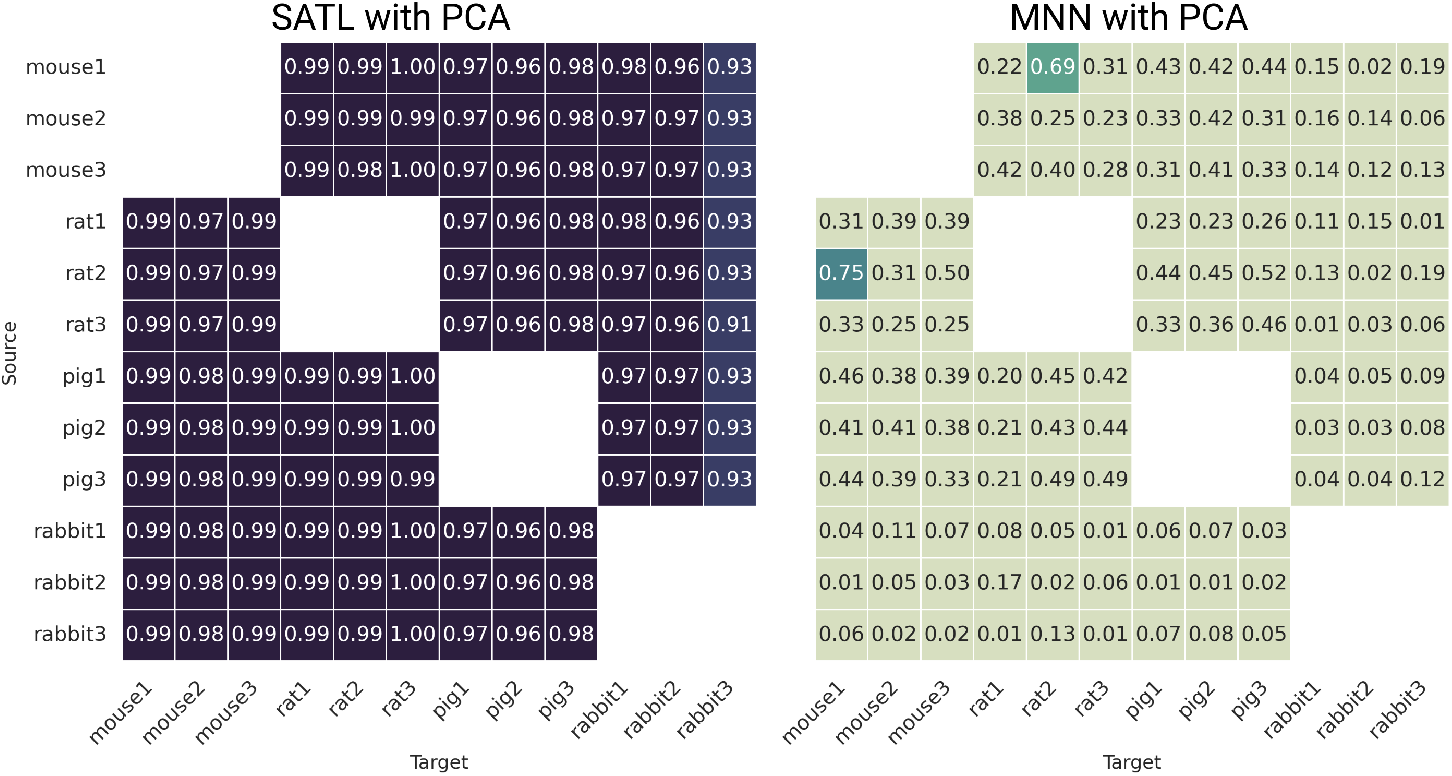
Comparison of out-of-sample label prediction with heterogeneous features. The xy-plot shows the results of the cross-species integration analysis comparing SATL with the MNN approach for heterogeneous domain adaptation. For evaluation, each RF model is trained on the source and tested on the target data with a masked label across species.

### 2.3 SATL prediction models identify aligned latent spaces with similar biological function

As shown in section 2.2 SATL models allow for the prediction of unseen classes in cell type labels across species. In order to evaluate the biological context of the aligned latent feature spaces derived by SATL based on the principal components of the single-cell RNA sequencing data obtained from different species, we performed a functional analysis using the gene ontology-biological processes [17], see Methods for details. We exemplarily examined the common latent spaces of pig and rabbit as shown in Figure 3. Figure 3a shows the aligned latent spaces of the prediction scenario where the *Pig1* dataset was used as source species and Rabbit 1, 2 and 3 are each used as target species (’Pig1 *→* Rabbit 1,2,3’) while ‘lps4’ cells were masked and predicted for each target dataset. One can see that masked cells from the rabbit data are correctly predicted based on the pig data on the common latent space. To validate whether SATL can find biologically plausible common latent spaces from all genes, we further investigated the variable importance. Therefore, the explained variance from PCA and the weight of the projection matrix from SATL are multiplied to find the most influential genes on each axis of the latent space.

**Fig. 3.**
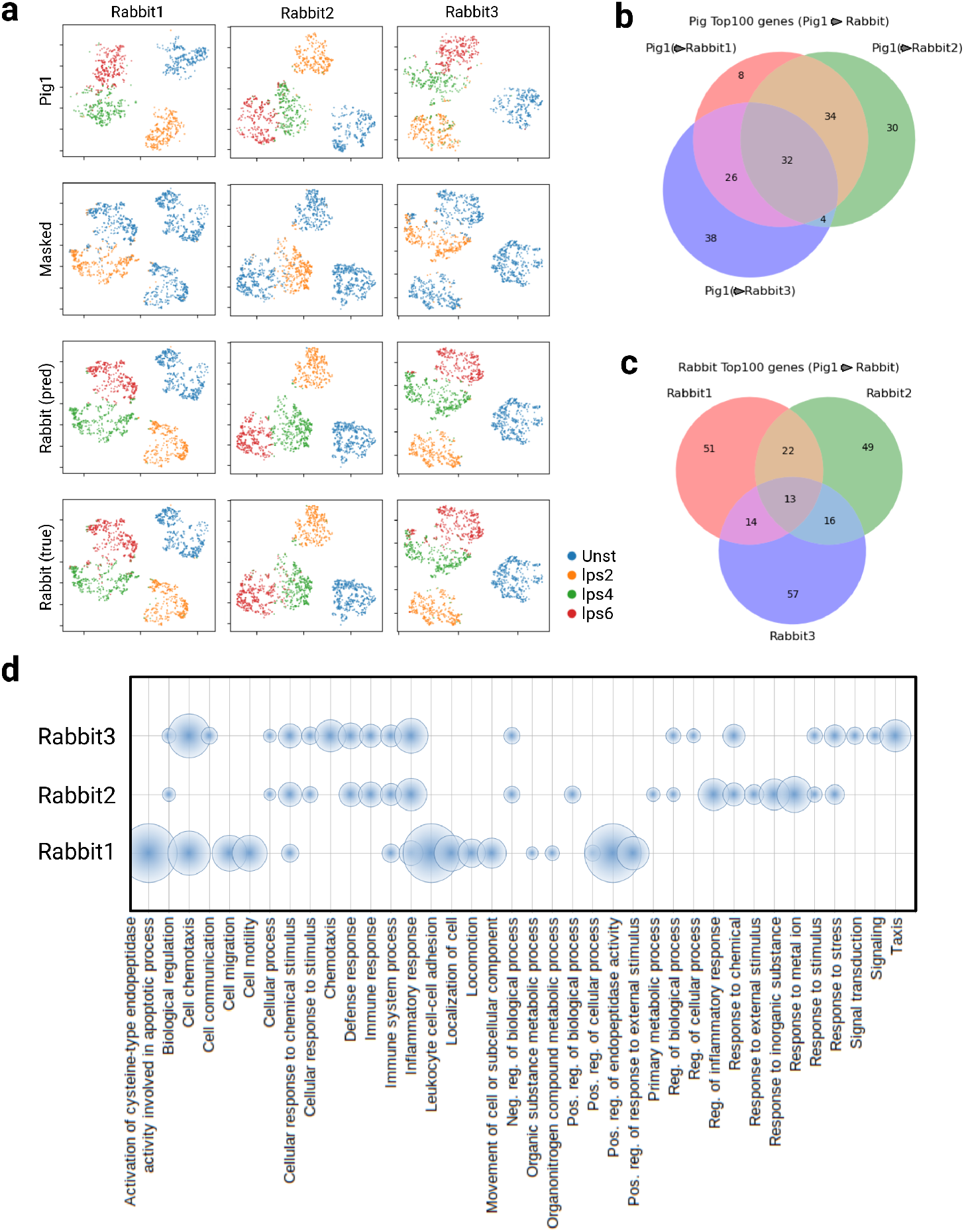
Visualization and interpretation of the common latent space. Shown is an example between pig as source and rabbit 1-3 as target in the lps-stimulated cell dataset. (A) Visualization of the *Source True Label (Pig)*, *Masked Cells (Rabbit)*, *Target Prediction (Rabbit)*, and *Target True Label (Rabbit)* in corresponding latent spaces: *pig*1 *→ rabbit*1, *pig*1 *→ rabbit*2 and *pit*1 *→ rabbit*3. SATL-aligned latent spaces are visualized via t-SNE plot. It shows the location of cells in the common latent space. (B,C) Overlap of 100 most important genes comprising latent space in Pig1 and Rabbit 1-3 depicted as Venn diagrams. (B) Pig1’s most important latent space genes aligned with different rabbit batches. (C) Rabbit1 3’s most important latent space genes aligned with Pig1. (D) Gene ontology (Biological Process) analysis of the 100 most important contributing genes on latent axis 1 from Rabbit1-3. Top enriched GO biological processes from ShinyGO are aggregated and shown. The radius of a circle means Fold Enrichment.

The top 100 genes from the first axis of *Pig1* and the *Rabbit1,2 and 3* data were examined. By comparing the later gene sets, we could observe a large within-species inter-individual variability in the latent space representation. Each rabbit sample resulted in a different common latent space when aligned with ‘Pig1’ data (Figure 3a). In particular, when investigating the top 100 important genes of the first axis of the common latent space, one can discover distinct sets of genes across the different biological replicates. This can be seen in Figures 3b and c depicting the venn diagrams of important genes composing the first axis of latent spaces in Pig and Rabbit, respectively. The venn diagram of ‘Pig1’ gene sets shows different features when they are integrated with different biological replicates of rabbits (Figure 3b,c). In the rabbit dataset, all three individuals share 13 genes out of the top 100 genes. In the paired pig1 result, when the same data is integrated with different rabbit individuals, they shared 32 genes out of the top 100 genes. These results can lead to individual variance or requirement for proper normalization steps in feature importance analysis with these SATL-aligned latent spaces. Furthermore, we examined the functional relevance of the top 100 genes in the rabbits by *Gene Ontology* (GO) enrichment of biological processes using ShinyGO [17]. As expected, the variation in the biological replicates and their important gene sets results in non-identical GO enrichment. Nevertheless, it reveals a general tendency towards immune-related GO terms (Figure 3d). Moreover, the GO enrichment analysis shows all three rabbits share the following biological processes: ‘Cellular response to chemical stimulus’, ‘Immune system process’, and ‘Inflammatory response’, for instance. Rabbit two and three are further sharing immune response, such as, ‘Cellular response to stimulus’, ‘Immune response’, ‘Response to stress’, and ‘Defense response’. These terms are also associated with the immune system. In contrast, Rabbit one shows ‘Leukocyte cell-cell adhesion’, ‘Pos. reg. of endopeptidase activity’, and ‘Activation of cysteine-type endopeptidase activity involved in apoptotic process’. This finding shows that, although identified important genes were different, they have similarities in their biological function, for instance, the presented Rabbit samples share similar immune system related GO terms while Rabbit one has a distinct response compared to the others.

### 2.4 SATL outperformed existing GZSL model and DA method in cross-species integrative analysis on three single-cell datasets

Finally, we performed a thorough comparison of SATL and related machine learning methods with and without prior knowledge on single-cell sequencing data from paired mouse and human organs, bone marrow, pancreas, and brain. Each dataset comprises different cell types and numbers thereof, see Methods 4.2. Similar to the previous analysis, we excluded the randomly selected two labels (here cell types) during the training process and predicted them in the target domain. Therefore, we combined SATL with three of the most common preprocessing strategies for single-cell sequencing data, namely Seurat (with PCA), DCA [18], and scETM [7]. Seurat with PCA is a common single-cell data analysis method, including a preprocessing step for filtering highly variable genes. Typically, the top 2,000 genes are selected for analysis. DCA additionally includes a step for filtering variable genes after its denoising process. To evaluate a diverse set of preprocessing strategies, we extracted features using scETM without a variable gene filtering step. Subsequently, we applied SATL to identify a common latent space between the mouse and human datasets, see Methods for details.

We compared our SATL results to CADA-VAE [19] which is a well-known method frequently showing good overall performance in generalized zero-shot learning image classification tasks. In addition to the default CADA-VAE model, we added CADA-VAE2, which employs two different VAEs for each species. Furthermore, we applied Quasi-Fully Supervised Learning (QFSL), a popular generalized zero-shot learning method [20]. However, for QFSL small changes in either the weight of the regularization or the classification loss led to collapsing representations on a shared space, in which seen classes are well separable, while the model completely fails to predict unseen classes, see Table 1. Moreover, we compared SATL to recent semi-supervised HDA methods, such as LPJT [21] and DDA [22]. Finally, we compared our results with recent methods that use prior knowledge to map data into the same feature space. Both scNym [23] and scAdapt [5] work completely unsupervised but utilize prior knowledge of orthologous genes. Both HDA methods were evaluated in a GZSL setting and a semi-supervised setting.

**Table 1.** Comparison of SATL, combined with different feature extraction methods (Seurat, DCA and scETM), with other state-of-the-art zero-shot learning and HDA approaches, such as CADA-VAE, CADA-VAE2, QFSL, LPJT and the gene orthology based methods scAdapt and scNym. Evaluation is based on the balanced accuracy of predicted unseen classes.

|  | BM | Pancreas | Brain |
| --- | --- | --- | --- |
| *Seurat*:SATL | 0.427 | 0.531 | 0.646 |
| *DCA*:SATL | 0.417 | 0.403 | 0.617 |
| *scETM*:SATL | **0.726** | **0.615** | **0.822** |
| *Seurat*:CADA-VAE | 0.120 | 0.212 | 0.322 |
| *DCA*:CADA-VAE | 0.139 | 0.177 | 0.194 |
| *scETM*:CADA-VAE | 0.201 | 0.132 | 0.323 |
| *Seurat*:CADA-VAE2 | 0.236 | 0.139 | 0.368 |
| *DCA*:CADA-VAE2 | 0.206 | 0.208 | 0.243 |
| *scETM*:CADA-VAE2 | 0.212 | 0.106 | 0.361 |
| *scETM*:QFSL | 0.243 | 0.116 | 0.228 |
| *Seurat*:LPJT | 0.066 | 0.050 | 0.099 |
| *DCA*:LPJT | 0.116 | 0.068 | 0.119 |
| *scETM*:LPJT | 0.140 | 0.064 | 0.100 |
| scAdapt* | 0.734 | 0.924 | 0.952 |
| scNym* | 0.780 | 0.538 | 0.623 |

\* Methods are tasks with prior knowledge.
BM: Bone marrow dataset

Table 1 presents the balanced accuracy of unseen classes across all three datasets. Since the balanced accuracy of seen classes was good for all methods, we focus on the balanced accuracy of unseen classes. Despite the different feature sets in paired species, SATL reasonably predicts the unseen cell types in the target domain. The performance varies depending on the dataset and feature extraction method used. However, scETM, which does not apply highly variable gene selection, achieves the highest balanced accuracy on all three datasets. SATL outperforms all other methods without prior knowledge. Although CADA-VAE2 shows slightly better overall performance than CADA-VAE, neither can beat SATL in terms of unseen class accuracy. Since HDA and LPJT are not designed for the GZSL task, they failed to predict unseen classes in the GZSL setting. Although LPJT has a similar concept with CDSPP by preserving the local structure, LPJT shows poor performance on the GZSL task. This can be explained by the fact that LPJT does not utilize labels of target data during the training. However, both methods show good performance in the semi-supervised task (Table B6 in Appendix). As expected, the overall accuracy of scAdapt, which uses prior knowledge, is better than the accuracy of the SATL and CADA-VAE methods. However, it should be noted that this is not a fair comparison as we are comparing a classification task with prior knowledge to a generalized zero-shot learning classification task. Nevertheless, this result demonstrates the possibility of directly integrating different species datasets without external knowledge.

In order to clarify the necessitate of the HDA method in dealing with heterogeneous features, we compared homogeneous domain adaptation methods using heterogeneous features obtained from PCA with a dimensionality of 50. Compared to heterogeneous domain adaptation methods like SATL, general domain adaptation methods developed for a homogeneous task cannot predict out-of-sample cell-type labels (Table 2). During our analysis, many domain adaptation methods collapsed and were not able to predict even seen classes. IWC, FA, PRED, KMM, and TrAdaBoost were able to predict seen classes reasonably. However, it failed to predict unseen classes. Thus no domain adaptation method outperformed SATL to enable transfer learning across different species with features extracted from PCA(n=50).

**Table 2.**
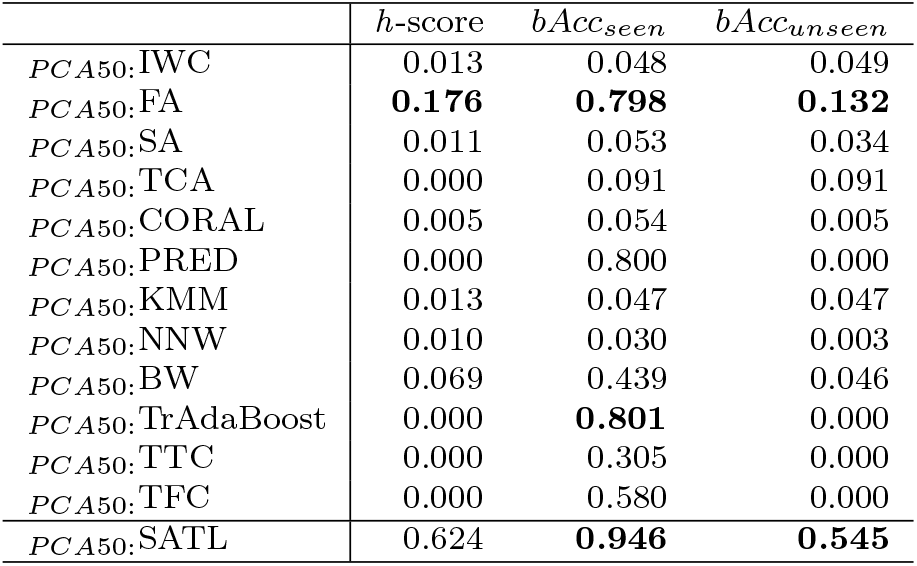
Comparison of SATL with homogeneous domain adaptation methods evaluating the out-of-sample prediction performance (h-score and balanced accuracy of seen and unseen classes) on the single-cell sequencing pancreas dataset (Mouse to Human)

### 2.5 SATL captures biologically meaningful latent space from mouse to human

To further validate our approach we utilized another dataset consisting of single-cell sequencing data from paired mouse and human organs, bone marrow, pancreas, and brain. Each dataset comprises different cell types and numbers thereof, see Methods 4.2. Similar to the previous analysis, we excluded the randomly selected two labels (here cell types) during the training process and predicted them in the target domain. The overall steps of this SATL evaluation were as follows: First, we reduced the dimensionality and extracted features in an unsupervised manner, including preprocessing with Seurat and PCA, DCA [18], and scETM [7]. Subsequently, we applied SATL to identify a common latent space between the mouse and human datasets, see Methods for details.

When investigating individual cell type-wise accuracy, we observed a drop in accuracy for specific cell types. Among the three datasets, the SATL model trained on the brain dataset performs best when cell labels are transferred from mouse to human (Table A3 in Appendix). This is likely due to the fact that the brain dataset shows balanced class distribution compared to the other two datasets. For the prediction of L2/3 IT cells, the scETM-SATL method shows slightly better performance than scAdapt, which uses external gene-pair knowledge. On closer inspection, the misclassified L2/3 IT cells were assigned to L5 IT cells (Figure 4). Literature search suggests that both cell types are closely related to intratelencephalic neurons and therefore share similar gene expression.

**Fig. 4.**
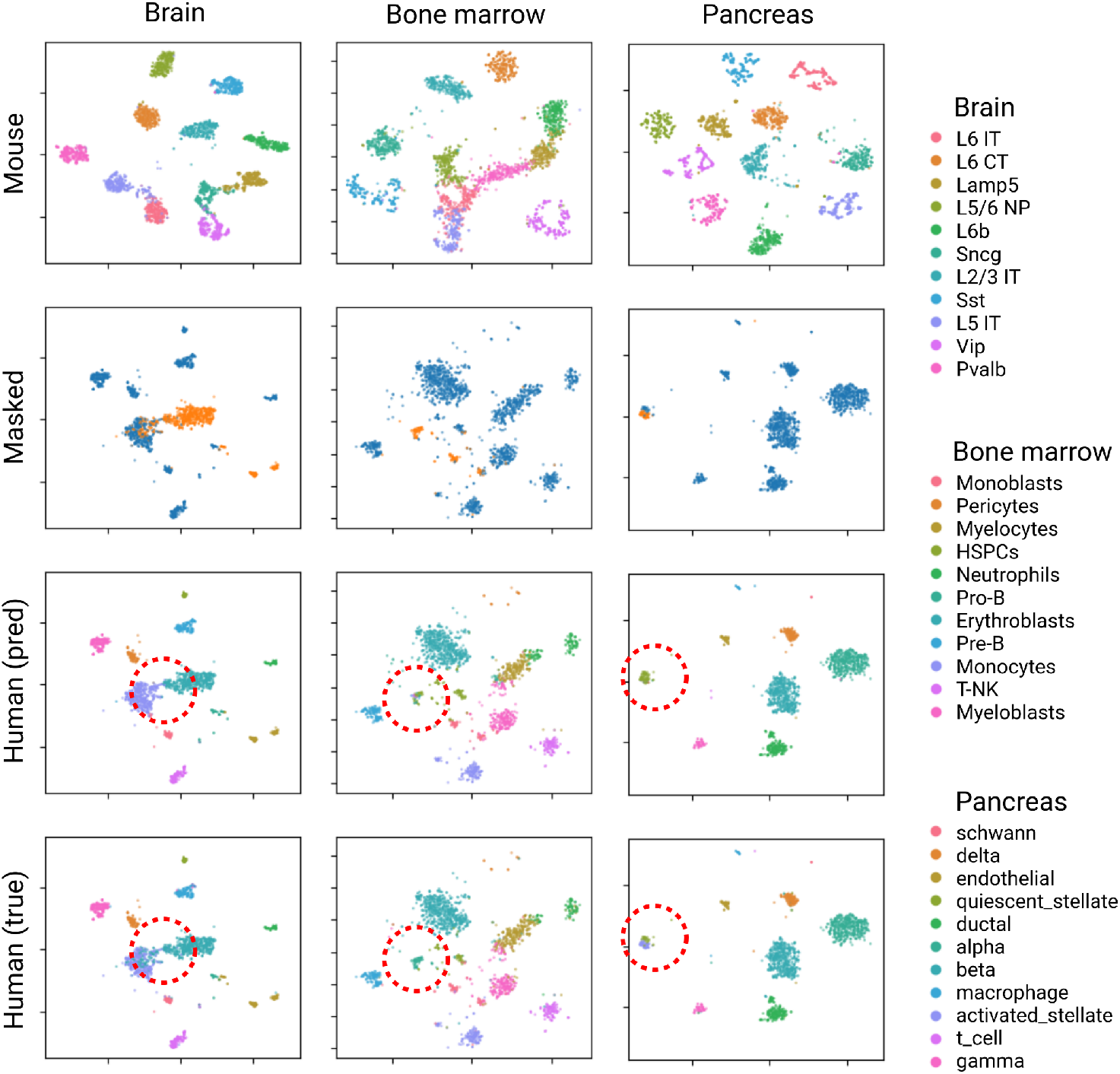
SATL latent space result representation using t-SNE from mouse to human cell-type label transfer analysis. scETM feature is used here. Masked cells are colored with yellow in the second row. Among masked cells, misclassified cells are highlighted with red circles in the third and fourth rows. In brain data, ‘Lamp5’ and ‘L2/3 IT’ cells are masked. ‘L2/3 IT’ cell is predicted to ‘L5 IT’ cell. In bone marrow data, ‘Monoblasts’ and ‘Pro-B’ cells are masked. ‘Pro-B’ is predicted to ‘HSPCs’. In pancreas data, ‘quiescent stellate’ and ‘activated stellate’ are confused.

In the bone marrow dataset, the accuracies of Pro-B and Pre-B cells were outliers among all other cell types (Table A1 in Appendix). The two cell types, Pre-B cells and Pro-B cells, are hard to distinguish in the latent space, resulting in Pre-B cells being confused with Pro-B cells and Pro-B cells being classified as HSPC cells. When Monoblasts and Pro-B are masked Pro-B is predicted to HSPC ((Figure 4). While this is a clear limitation of our model, it can be explained by looking at the cell differentiation lineage tree, where these cell types are closely related cell states. Specifically, Pro-B cells are on a developmental level located between Pre-B and HSPC cells. However, this differentiation was also challenging for the method using external knowledge, scAdapt (Table A1 in Appendix).

In the pancreas dataset, several cell types are rare in the mouse dataset, namely t-cells, schwann, activated stellate cells, and macrophages. Consequently, all methods, SATL, as well as methods with external knowledge show difficulties identifying these cell types (Table A2 in Appendix). While the performance is relatively stable with the other preprocessing techniques, in combination with scETM-features, the model fails to recognize two out of these four cells. When ‘activated stellate’ is masked, it is predicted to ‘quiescent stellate (Figure 4). A second reason for this bad performance is that there are different degrees of biological similarity. When studying the correlations between the average expression profiles of these cell types, it becomes clear that macrophages, T and Schwann cells are less correlated between different species [24], further increasing the difficulty of zero-shot classification with species agnostic transfer learning.

These examples demonstrate that the generalized zero-shot classification errors of our approach are not simply random, but rather are based on biological reasons. However, at the same time, the results showing different performances on different features highlight the importance of the feature extraction method for SATL analysis. Since HDA depends on the extracted features, if the feature extractor fails to embed an appropriate gene signature that can classify similar cell types, the performance of SATL will be limited by the applied feature extraction methods.

## 3 Discussion

In this paper, we present SATL, a novel methodology for species-agnostic transfer learning with heterogeneous domain adaptation in biomedical research. To the best of our knowledge, SATL is the first algorithm that allows cross-species data integration without relying on any prior knowledge such as gene homology information. The CDSPP, baseline model of SATL, solves a heterogeneous domain adaptation using a latent embedding approach with a supervised eigenvalue problem by utilizing seen-label information. This CDSPP-based method can have a significant advantage in computational efficiency compared to recent deep neural network models which require training for numerous parameters. In contrast to other methods, the key idea of SATL is to utilize latent feature vectors for domain adaptation instead of gene-expression profiles. This characteristic enables a versatile integration of various species datasets using different feature extraction methods on gene-expression profiles while minimizing information loss which is a consequence of gene-id conversion. The feature, latent-feature-based integration, gives additional benefit to our model. This SATL can integrate any deep features extracted from various high-throughput data, such as transcriptome, genome, epigenome, proteome, or metabolome. This means SATL shows its possibility on cross-species data integration, however, it is possible to integrate multimodal data for a model organism study. Nevertheless, there are also disadvantages to the suggested approach. As mentioned in the original CDSPP study and shown in Table 1, this SATL approach also relies on the performance of the feature extractor. Another drawback of SATL is its limitation towards applications in the transductive generalized zero-shot learning setup, which requires both labeled and unlabeled test samples to improve the accuracy of recognizing unseen classes. Additionally, we have constrained our analysis to situations where both domains have the exact same set of classes, meaning that the two domains have the same set of cell labels. This constraint can limit the ability to identify novel cell types in cross-species studies. In the next step, we could extend the analysis with a dataset having highly-imbalanced classes.

The proposed training and testing scheme for cross-species transfer learning in GZSL application is easily expanded and applied to various biomedical research problems. SATL learns a projection matrix for common latent space to handle the GZSL task where partial common data with labels are available. Compared to the unsupervised/semi-supervised task, this GZSL task has great benefit in expanding knowledge based on accumulated datasets in biomedical domains. Our first analysis on the lps-stimulated dataset with the four species dataset demonstrates that SATL learns a biologically meaningful latent space and can predict unknown cell types through this latent space. The projection further enables the observation that masked lps4-cells are located between lps2 and lps6 cells. This result shows the possibility of cross-species-wise RNA velocity work with the aligned latent space identified by SATL. Another interesting aspect is intra-species individual variability while the biological replicates, rabbit1, rabbit2, and rabbit3, share a similar GO term related to ‘innate immune response’ which is the target biological process the original study designed.

As described, the SATL approach has an advantage in integrating all features in the data. When evaluating the ten most important genes in the first axis of the latent SATL space of Pig one, two genes, namely ENSSSCG00000028525 and ENSSSCG00000000246, have no match in databases (Not found in PantherDB 17.0 [25]; Retired in Ensembl release 110-July 2023). However, further analysis with Fold-seek [26] detected two structurally very similar proteins (Figure 5). Moreover, these two proteins, *ENSSSCG00000028525-CXCL2* and *ENSSSCG00000000246-SAA2*, are reported to be immune-system-related [27, 28]. Moreover, when the ten most important genes of the latent space of Rabbit one are queried in the orthologue analysis database, Better Bunny (Aug. 2020) [29] and Ensembl (release 110-July 2023), ENSOCUG00000022364 is reported as a novel, yet unknown rabbit gene orthologues to interferon-induced transmembrane protein. Foldseek search found the same protein as a structurally similar protein. Those genes are dropped in a gene-id conversion preprocessing. However, in SATL analysis, they are found as important genes to investigate lps-stimulated response on macrophages. This result can illustrate a limitation of the current gene-id conversion method and a potential new approach to discover and investigate functional similarity of genes between different species.

**Fig. 5.**
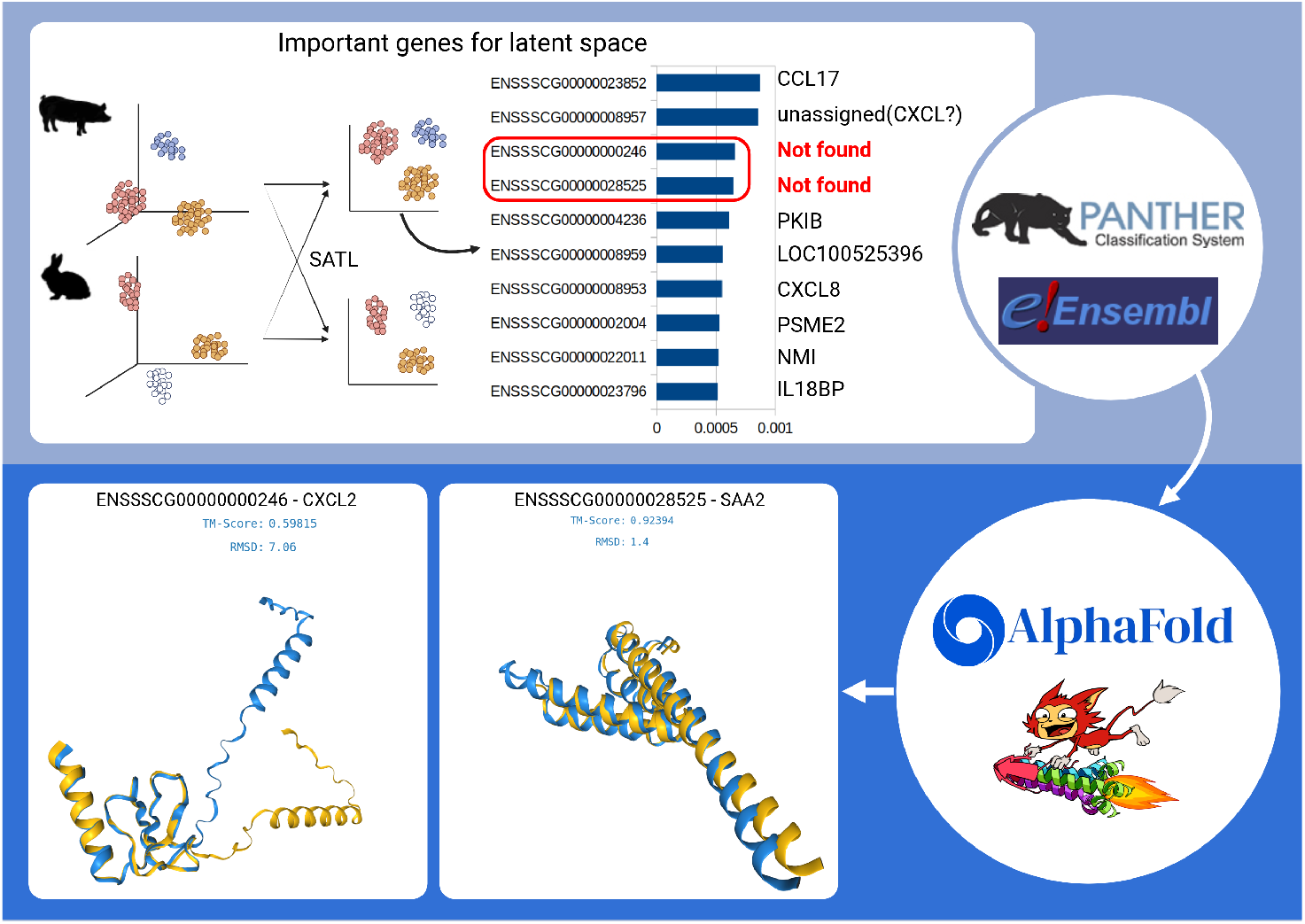
Phenotype-centric orthology inference via SATL. Proposed approach for ortholog analysis with SATL is shown. SATL exploits labels of data and finds latent space related to experimental design. When important genes composing latent space are investigated, some genes are not found in the current orthologous database. Two genes are retired in the Ensembl database. Therefore, structural similarity analysis of protein is done. Foldseek search identified ‘ENSSSCG00000028525’ as a similar protein to CXCL2 with sequence identity: 75.7 and E-Value: 3.87e-14. ‘ENSSSCG00000000246’ is identified as a similar protein to SAA2 with sequence identity: 72.3 and E-Value: 2.47e-13.

**Fig. 6.**
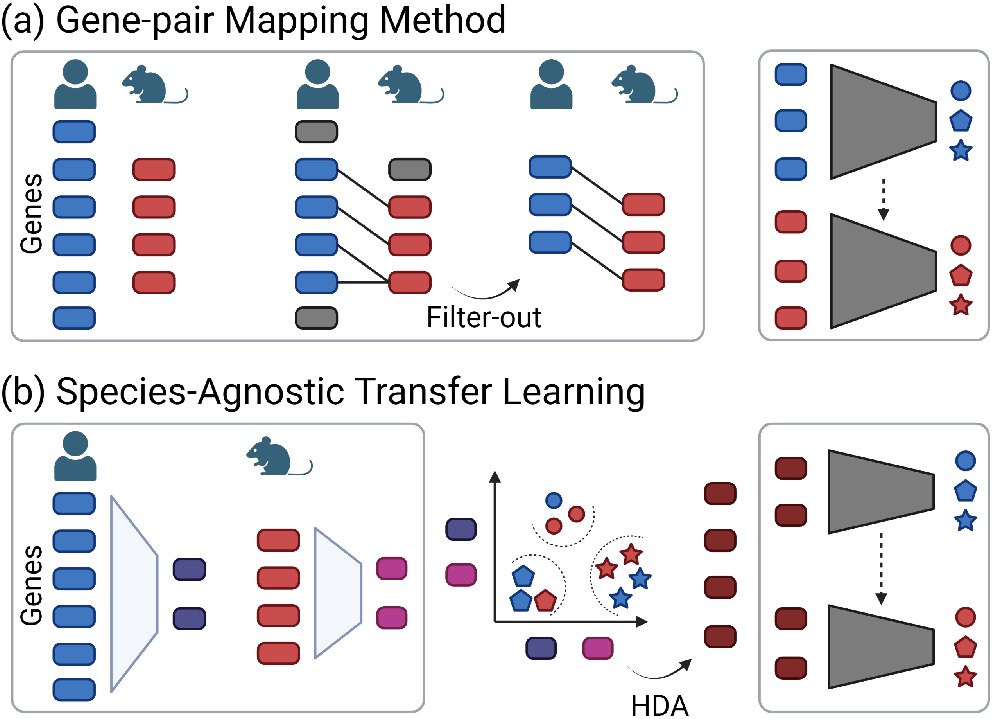
Comparing the strategy of previous approaches with the newly developed SATL method. (a) Previously, transfer learning across species data is done using external gene homology data. However, this approach often results in severe information loss. (b) In the proposed SATL method, a feature extractor is used to embed a gene set into latent features. After feature extraction, heterogeneous features are aligned using a heterogeneous domain adaptation (HDA) algorithm. The aligned features can then be used directly in the integrative analysis.

In summary, we present a new data-driven method, SATL, to enable the integration of heterogeneous high-throughput data across various species. In particular, SATL shows great potential for heterogeneous domain adaptation algorithms to expand our species-dependent knowledge to more general pan-species biological knowledge. SATL can dissect functional groups of genes in the species-common latent space when presented with large-scale sequencing data, such as the responses of organisms under various experimental conditions. Current bioinformatics algorithms can predict orthologous genes based on sequence homology but cannot predict analogous genes with similar functions but dissimilar sequences. One of the alternative approaches is structure-based protein function prediction, such as AlphaFold [30]. A protein language model from a functional perspective can be a good alternative method for alignment-based orthologous or homologous gene information [31]. Still, we hope that SATL can aid further investigations of knowledge in various species on the evolutionary tree of life with a data-driven approach. Our approach can be beneficial as a new analogous gene prediction approach, phenotype-centric, in the era of big sequencing data.

## 4 Methods

### 4.1 Task details: out-of-sample label prediction with a latent feature from single-cell sequencing data

The learning and testing scheme follows a transductive GZSL setup where the labels of a certain fraction of classes in the target domains are unknown during training. The machine learning model has fully labeled data from a source domain. However, in the target domain, only a partial set of classes are labeled and available during the training step. The main task of this work is an out-of-sample prediction in a target species. Without gene-ID conversion, which incurs severe information loss, we first extract features and then integrate the features of different species to project labels from one species to another. We expect this new approach to have a better representation of the transcriptomic features of each species than a preprocessing step with highly-variable gene selection after gene-ID conversion. The classifier will be evaluated using target species samples from all classes, both known and unknown during training. By masking all different combinations of cell types and examining the results, this will provide insights into the possibilities of a common representation of different species with biological meaning.

In particular, for the task with sequencing data, the domain is the species, and the data is the gene expression profile. Domain heterogeneity comes from biological heterogeneity, as different species have different sets of genes, and the overlapping functions/genes between species vary depending on the evolutionary distance of species. The label projection with single-cell sequencing data is done by cell types (Bone Marrow, Brain, Pancreas dataset) or experimental labels (Stimulation dataset). Thus, the dataset of single-cell sequencing is composed of gene expression and corresponding cell labels (number of cells x number of genes & Cell Labels). The cell types are determined from the expression profile of a set of marker genes. Since different species have different gene set, the same cell types in different species can be determined by a different set of genes.

The general workflow is shown in Figure 1. First, in each species, the gene expression profile is converted into feature vectors. The feature extraction methods can be PCA, *Autoencoder* (AE)-based models, or any other available tools. Second, the feature vectors from each domain are integrated with heterogeneous domain adaptation methods. During this step, common cell labels from domains are incorporated and used to adjust two different feature spaces into a common latent space. Lastly, in the common latent space, it is possible to predict the labels of the target species based on the knowledge from the source species.

### 4.2 Dataset and Preprocessing

To show cross-species knowledge transfer, we analyze four datasets. Three datasets comprise single-cell sequencing data with cell labels of bone marrow [4], pancreas [24], and brain (https://portal.brain-map.org/) both from human and mouse. Each dataset has eleven different cell populations. The fourth dataset originates from a lps-stimulation experiment with four species, mouse, rat, rabbit, and pig (E-MTAB-6754) [16]. There are three biological replicates for each species, and each dataset has four different cell-differentiation statuses. The overall observation counts, as well as the number of genes and total cell counts, can be found in Table 3.

**Table 3.** Description of datasets after quality control. Datasets are used for the comparison of SATL performance in cell type label projection analysis.

| Tissue | Species | Total Cells | Genes | Cell types |
| --- | --- | --- | --- | --- |
| Bone marrow | Human | 9234 | 20887 | 11 |
| Bone marrow | Mouse | 5504 | 14909 | 11 |
| Pancreas | Human | 8567 | 15858 | 11 |
| Pancreas | Mouse | 1883 | 13653 | 11 |
| Brain | Human | 9564 | 29499 | 11 |
| Brain | Mouse | 15396 | 22428 | 11 |

The analysis was performed in two different ways: using all cells in the original data without any preprocessing, and using preprocessing techniques such as over/under-sampling. If no preprocessing method was specified, the analysis was performed on the RAW data. The preprocessing involved several steps. First, cells with fewer than 200 expressed genes were discarded, as these are often of low quality due to their low gene capture rate. Only genes expressed in at least 5 cells were considered to reduce the number of features [32]. Cells with inflated gene counts, typically caused by doublets during sequencing [33], were also filtered out. In addition, cells with high counts for mitochondrial genes, indicative of perforated cells, were excluded from the analysis [34]. The gene count matrix is normalized with ‘Total-count with 1e6’ and log-transformed. The subsequent steps depended on the chosen method, which will be described in subsequent sections. Table 3 shows the preprocessed dataset used in this study. For the lps-stimulated cell dataset, preprocessing is done in the similar procedure. First, cells with fewer than 200 expressed genes were discarded, as these are often of low quality due to their low gene capture rate. Only genes expressed in at least 5 cells were considered to reduce the number of features. The gene count matrix is normalized with ‘Total-count=1e6’ and log-transformed. Table 4 shows the preprocessed dataset used in this study.

**Table 4.** Description of mononuclear phagocyte stimulation dataset used for SATL comparison with MNN and common latent space analysis. It shows the number of genes after single-cell preprocessing.

|  | mouse1 | mouse2 | mouse3 | rat1 | rat2 | rat3 |
| --- | --- | --- | --- | --- | --- | --- |
| #Gene | 12962 | 13070 | 12826 | 13350 | 12821 | 12613 |
| unst | 2126 | 3519 | 2936 | 2650 | 1748 | 1960 |
| lps2 | 2537 | 4321 | 3169 | 2094 | 2682 | 1962 |
| lps4 | 2366 | 3293 | 3536 | 2779 | 1718 | 1079 |
| lps6 | 1703 | 2126 | 1453 | 3909 | 1362 | 2526 |
|  | rabbit1 | rabbit2 | rabbit3 | pig1 | pig2 | pig3 |
| Genes | 10899 | 10694 | 10853 | 12041 | 11846 | 12121 |
| unst | 3773 | 3275 | 3399 | 2221 | 1748 | 2179 |
| lps2 | 3690 | 1820 | 4401 | 1630 | 1614 | 2353 |
| lps4 | 2965 | 1522 | 2163 | 1948 | 1899 | 2758 |
| lps6 | 4196 | 1660 | 1664 | 1941 | 1381 | 1797 |

**Table 5.**
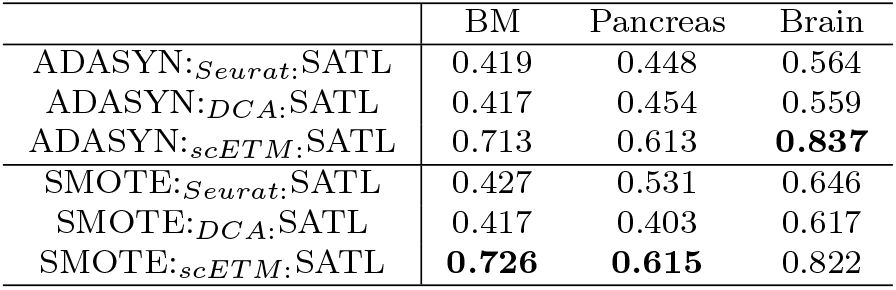
SMOTE and ADASYN comparison. The majority classes will be randomly under-sampled and synthetic samples for the majority classes created using SMOTE or ADASYN. The observed variations in performance are not significantly meaningful between SMOTE and ADASYN.

|  | BM | Pancreas | Brain |
| --- | --- | --- | --- |
| ADASYN:*Seurat*:SATL | 0.419 | 0.448 | 0.564 |
| ADASYN:*DCA*:SATL | 0.417 | 0.454 | 0.559 |
| ADASYN:*scETM*:SATL | 0.713 | 0.613 | **0.837** |
| SMOTE:*Seurat*:SATL | 0.427 | 0.531 | 0.646 |
| SMOTE:*DCA*:SATL | 0.417 | 0.403 | 0.617 |
| SMOTE:*scETM*:SATL | **0.726** | **0.615** | 0.822 |

The selection of preprocessing methods in this study was informed by recent standard analysis pipelines and encompassed both linear and non-linear feature extraction models. Specifically, we employed several methods, including the Seurat-pipeline [35] as the standard method, DCA [18] as a de-noising AE, and scETM [7] for deep feature extractor. These methods were chosen due to their established efficacy in de-noising and feature extraction. The scripts used for each preprocessing and feature extraction can be found in the codebase.

Seurat’s data preprocessing workflow involves several steps. To ensure comparability of expression counts across cells, counts are transformed into CPM-values. Next, the issue of heteroscedasticity is addressed by transforming counts using the natural logarithm. To reduce the dimensionality of the data, the number of genes is typically reduced to the 1000-5000 most variable genes. In this study, the top 2000 most variable genes are selected and divided into 20 bins based on their mean expression levels. Within each bin, the variances are z-transformed, and the 2000 genes with the highest transformed dispersion measures are retained for analysis. To remove the effects of absolute counts for each cell and mitochondrial counts, they are regressed out, and each gene is scaled to unit variance. To obtain low-dimensional representations, the data is reduced to the 50 most important principal components. This number was chosen since very low-dimensional representations can distort distances between samples, and the scETM [7] method also extracts a 50-dimensional embedding.

DCA is applied after quality control, and it internally applies further normalization. By running DCA on default values with three hidden layers (64, 32, and 64 units), a batch size of 32, and batch normalization, we obtain a denoised and imputed expression matrix. After applying DCA, we apply count normalization, natural logarithm transformation, reduction to highly variable genes, z-transformation, and PCA-based dimensionality reduction, similar to the Seurat preprocessing pipeline.

scETM is used as another unsupervised method that does not require preprocessing beyond quality control. It directly extracts low-dimensional features without reconstructing de-noised count data. By running scETM on default values with at least 6000 gradient steps, we obtain a 50-dimensional cell-topic embedding, which is used for the generalized zero-shot domain adaptation task. We chose scETM, a state-of-the-art unsupervised model that extracts features that have been shown to be useful for cell identification and transfer learning.

### 4.3 SATL details

The baseline model is a state-of-the-art heterogeneous domain adaptation method, CDSPP [36], which is the heterogeneous domain version of the Supervised Locality Preserving Projection (LPP) [37]. The objective of LPP can be transformed into:

Let ***P*** *_s_*and ***P*** *_t_*are the domain-specific projectors with *d^s^*, *d^t^*, *d^c^*being the dimensions of the source domain (***D****^s^*), target domain (***D****^t^*), and common latent space (***D****^c^*).

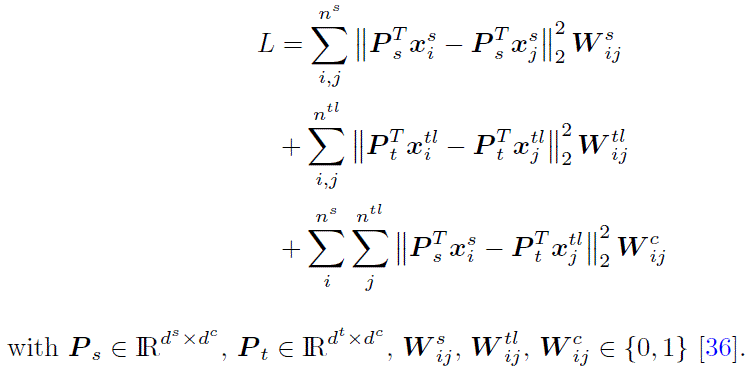

This structure-preserving transformation approach does not force class separability or minimize maximum mean discrepancy but preserves the original structure better [38]. In the same manner with CDSPP, at first, all observations are l2-normalized. The feature vectors, therefore, all have the same length. Based on the normalized features, the generalized eigenvalue problem is solved, and optimal projection matrix ***P*** *_s_* and ***P*** *_t_* can be obtained (details in [36]).

Since the goal of the task is a GZSL application and not a classical semi-supervised DA task, preserving the local structures of both domains is a crucial factor that can make a model avoid overfitting to seen classes and underperforming on unseen target classes. In summary, all data is then projected to the common latent space, centralized, and again normalized. After that, to obtain the class prototypes, the observations belonging to each class are averaged to calculate the coordinate of the prototypes. The out-of-sample prediction (*y_i_*of ***D****^u^*) is made by applying the same transformations to the test data, including classes both seen and unseen in the target domain for the GZSL task. Prototypes of classes not in the source domain are only calculated based on source domain data. Finally, the test data is classified according to the closest prototype.

To further reduce domain bias, various works propose the application of pseudo-labeling algorithms [36, 39]. In order to deal with both unseen classes and handle the imbalance of classes in the test set, the following pseudo-labeling scheme is proposed here 1. The algorithm is split into two procedures. In the first part, only predictions for out-of-sample classes are considered. Then, in each iteration, an increasing amount of test data with their respective prediction is added to the training data. In the second step, the process is repeated on all class predictions. This procedure considers that the initial model is strongly biased toward the source domain in the zero-shot setting. Therefore, we use an initial burn-in phase for out-of-sample classes. For this work, both parameters are set as *K* = 10 and *K_oos_* = 10.

In addition to that, due to the biological dataset consists highly imbalanced classes, we applied SMOTE [40] to oversample the minority classes in the main result (Table 1). This set-up follows the work by [41] where oversampling was applied after dimensionality reduction.

#### 4.3.1 Sampling methods for data balancing

Benchmark datasets for domain adaptation or generalized zero-shot learning are often well balanced, with similar numbers of samples per class in both training and test sets [10]. However, in real-world applications, this is often not the case, and all datasets have strongly imbalanced cell counts. As a result, biased models and estimators may be produced, presenting a consequential danger [42]. To handle this issue, three strategies are proposed, namely preprocessing (re-sampling), algorithmic-centered (reweighing the model loss), and hybrid approaches. This study employs a sampling method to balance the training data for both source and target domains. Specifically, the study adopts a combined approach of over-and under-sampling, where several approaches for each type of sampling, as well as their combinations, are available in the literature. The balancing is applied after the preprocessing procedure, and since the majority of classes in all data sets have around 300 observations, the size per class is set to 300.

**Algorithm 1.**
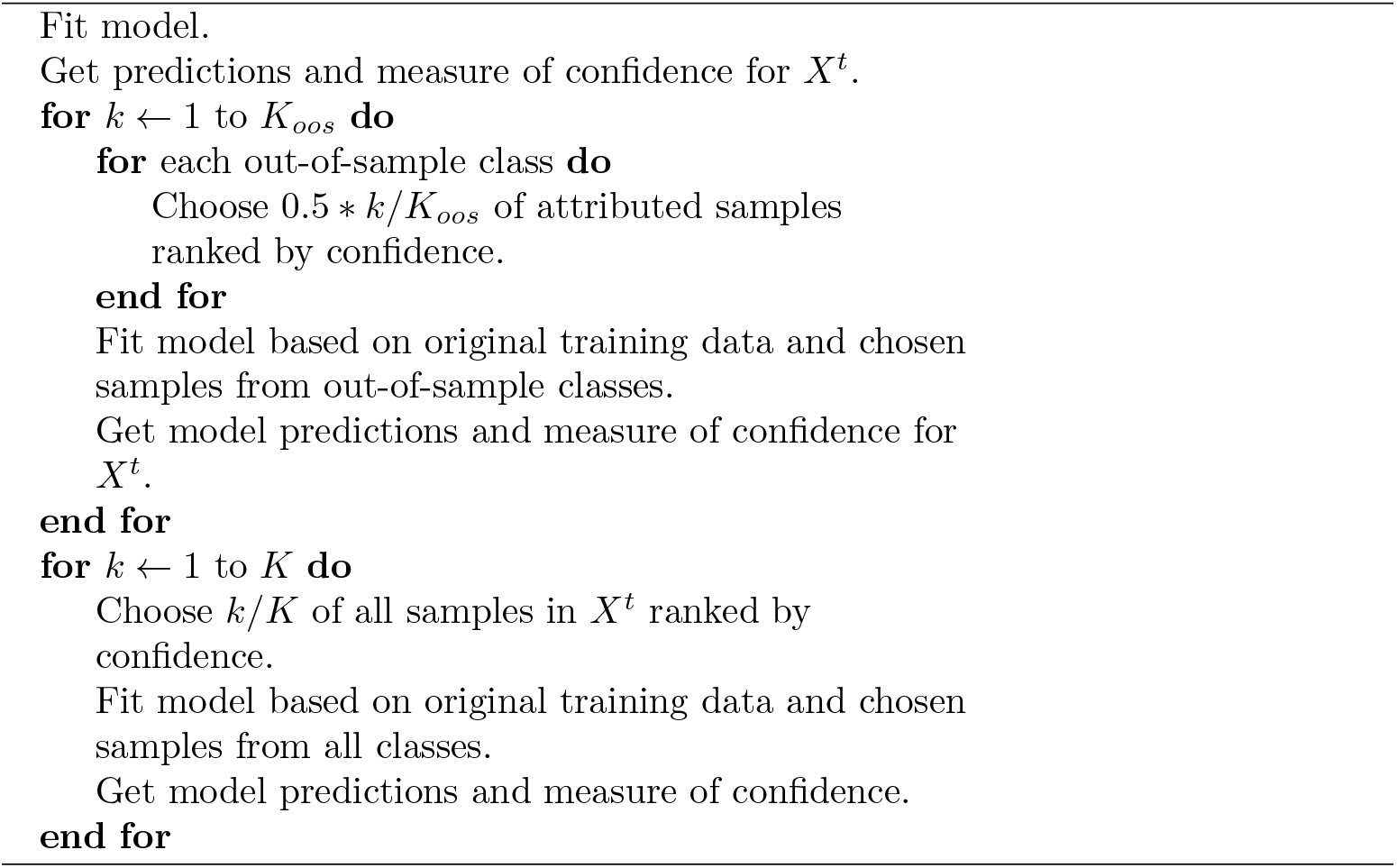
Pseudo-labeling for GZSL algorithm.

SMOTE [40] was applied to the main results. Random under-sampling was used for the majority classes, while over-sampling was used for the minority classes. The reason for this approach is that the data set is highly imbalanced (up to two orders of magnitude), making a combined approach the only feasible option [43]. SMOTE creates synthetic samples by generating them at random points along the lines that connect samples of the minority class with their k-nearest-neighbors.

ADASYN [44] works in a similar way to SMOTE by generating synthetic samples on the lines connecting the nearest neighbors of the same class. However, unlike SMOTE, the number of synthetic samples generated is set proportional to the number of neighbors of the same class in the kNN neighborhood. Therefore, if a sample has no other class member in its k-vicinity, no synthetic sample will be generated. Conversely, if all of a sample’s k-nearest neighbors are of the same class, the maximum number of synthetic samples will be generated on the lines connecting the neighbors. As such, in this project, the majority of classes will be randomly under-sampled, and synthetic samples for the majority classes will be created.

#### 4.3.2 Pseudo-labeling

In general, the pseudo-labeling approach with label projection involves several steps. In the initial run, a fraction of the test samples with the highest confidence or smallest distance to the class center are added to the training data with their predicted label. This partial update is repeated K times by increasing the ratio of pseudo-labeled test data by one (1/K, 2/K, …, K/K = 1). Although this approach has shown good performance in few-shot settings with very limited target domain data, it has a limitation when it comes to domain adaptation with generalized zero-shot tasks. The algorithm tends to be biased towards seen classes, as the confidence for seen classes is always higher than for unseen classes. Consequently, the model becomes more favorable to seen classes and assigns only seen class labels. To address this issue, an alternative version [45] has been developed. This pseudo-labeling scheme separates seen and unseen classes during the pseudo-labeling step, meaning that even if the prediction is sub-optimal in terms of confidence score, it could still be assigned to unseen classes. Another difference in [45] is how unlabelled data is integrated into the training set during each iteration. They keep the test samples that were chosen based on confidence as part of the training set for the whole pseudo-labeling procedure, and consequently, the labels will not be updated once they are accepted with the highest confidence prediction. While this approach is promising for better prediction of unseen classes, it is important to note that the method by [45] assumes a balanced test set.

In an unbalanced scenario, the proposed algorithm will assign all samples contributed to a minority class to the training set during the first iteration, risking bias based on predictions made by a model not yet calibrated. Additionally, once a prediction is accepted, there is no way to correct it afterward, even if the prediction would change during calibration. Furthermore, the results presented in the CDSPP work showed that while the h-score improves, the accuracy for seen classes decreases in half of the experiments [36]. This issue might become even worse in an unbalanced test setting.

The proposed algorithm in this work combines aspects of [45] and [36]. It consists of two phases. In the first phase, only predictions for unseen classes are considered for label projection to allow the model to consider more unseen classes. The amount of high-confidence samples chosen for each class is relative to the overall amount of samples attributed to that class. The parameter *λ* regulates the share of unseen class predictions to be considered as a labeled training set in the next iteration. If *λ <* 1, not all unseen class predictions will be considered as a training set to prevent a complete shift to unseen classes. Additionally, high-confidence samples are only temporarily added to the training set to allow the model to update its predictions of accepted test data. The labels of all test data are re-evaluated after each round of calibration.

#### 4.3.3 Dimensionality of latent features affects performance of SATL

To investigate the extent of information loss and the performance of SATL, we conducted experiments to test the model by varying feature sizes without applying a variable gene selection and oversampling method. We first tested the number of components of PCA from 2 to 800 and examined the explained variance on each component’s size while the number of latent space for SATL is eleven. The results showed that the balanced accuracy of seen classes reached over 90% at 50 components (Table 6). At this point, the cumulative explained variances were 99% and 99.6% in the human and mouse datasets, respectively. This indicates that features extracted from PCA and aligned with SATL can be reversed into the original gene expression profile with less than 1% of information loss (Table 6). In the PCA-SATL scenario, the balanced accuracy of unseen classes reached 50% while increasing the component size to 800. However, in the case of CADA-VAE2, although the accuracy of seen classes increased as the latent vector size increased, the accuracy of unseen classes did not change (Table 7).

**Table 6.** Evaluation of the influence of varying PCA dimensionality reduction on the SATL out-of-sample prediction performance and cumulative explained variation in the human and mouse pancreas RAW dataset. H-score and balanced accuracy of seen and unseen classes are shown. Scores are the averaged value of 10 repetitions. SATL without oversampling and pseudo-labeling are analyzed.

| PCA | Cumulative explained variances |  | Score |  |  |
| --- | --- | --- | --- | --- | --- |
| | Human | Mouse | $h$-score | $bAcc_{seen}$ | $bAcc_{unseen}$ |
| $n=2$ | 0.788 | 0.855 | 0.061 | 0.337 | 0.081 |
| $n=11$ | 0.976 | 0.992 | 0.402 | 0.793 | 0.337 |
| $n=50$ | 0.990 | 0.996 | 0.435 | 0.928 | 0.340 |
| $n=100$ | 0.993 | 0.997 | 0.529 | 0.951 | 0.423 |
| $n=200$ | 0.995 | 0.998 | 0.598 | 0.948 | 0.501 |
| $n=400$ | 0.996 | 0.999 | 0.582 | 0.963 | 0.489 |
| $n=800$ | 0.997 | 0.999 | 0.580 | 0.957 | 0.490 |

**Table 7.**
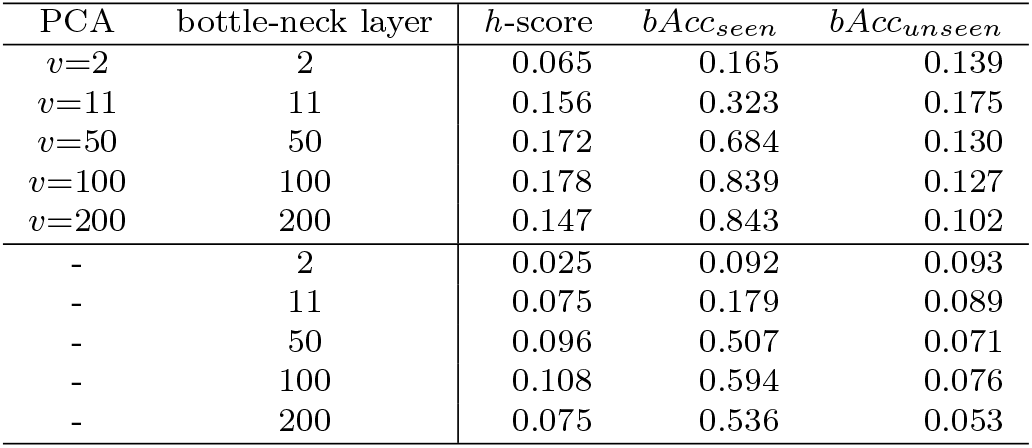
Comparison of different sizes of latent vectors generated by CADA-VAE2 on the mouse-human pancreas dataset. PCA and size of bottle-neck layers are shown. H-score and balanced accuracy of seen and unseen classes are shown. Scores are the averaged value of 10 repetitions.

Afterward, we tested the latent dimensionality parameter for SATL instead of fixing it at eleven. For this test, we used a fixed dataset from three different feature extraction methods. The results indicate that a small number of dimensions for SATL can distort the latent space and degrade its performance (see Figure 7). This is a similar finding to the original CDSPP work with the image classification task. In addition to that, a latent space with too large of dimensionality cannot significantly improve its performance. In most cases, a high dimensionality compared to the total number of classes to be addressed can diminish the performance of SATL.

**Fig. 7.**
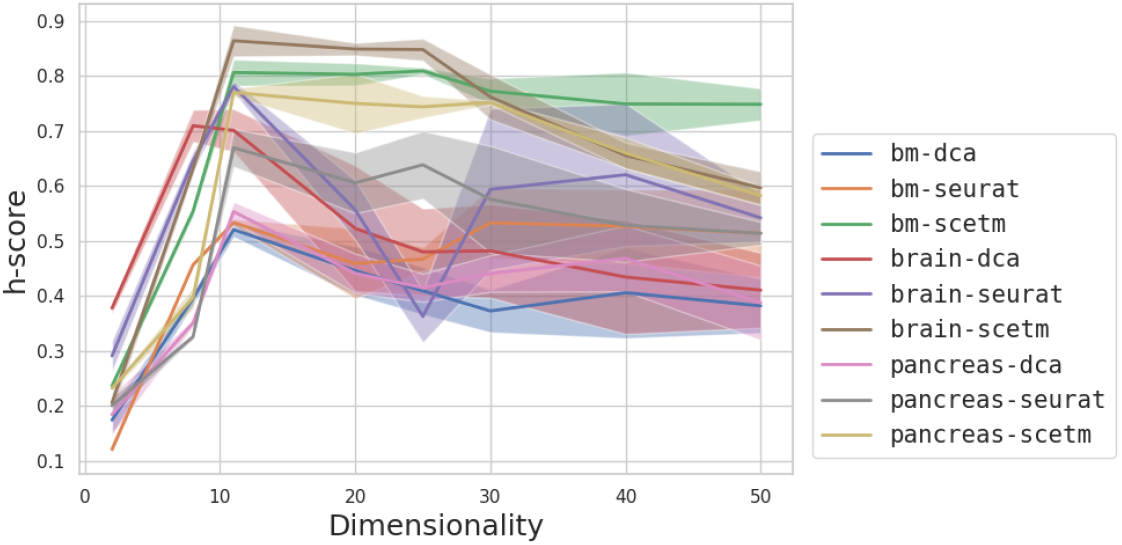
Visualization of the impact of dimensionality of the hidden layer on prediction performance. The shown h-score is evaluated for three different datasets and three different preprocessing and dimensionality reduction methods.

The use of human-mouse orthologous gene information for gene filtering typically results in the exclusion of tens of thousands of genes out of the nearly 20,000 total genes. However, in the recent work by [4], they filtered genes based on highly variable-geneset and orthologous genes information, resulting in the use of only 4,372 genes in the inter-species analysis. This information loss during gene conversion becomes more severe when less investigated species are incorporated into the analysis. However, compared to the previous method, the degree of information loss of the SATL method becomes negligible when the feature extraction method utilizes the whole gene expression profile.

SATL hyperparameter for numbers of dimensions is evaluated with the number ranging from 2 to 50 since preprocessed data have 50-dimensional features. In most cases, the overall h-score reached its highest value at dimensionality 11 or 20. As all three datasets contained eleven cell types, having a latent dimension that is too large for the number of classes may deteriorate SATL’s performance. An outlier was observed in the brain dataset when using the Seurat feature. For the comparison, we conducted a similar dimensionality test with CADA-VAE, a deep neural network model for generalized zero-shot learning tasks. We tested different dimensions of CADA-VAE in two ways: with PCA and without PCA. When we used PCA for feature extraction, the feature vector from the PCA transformation was fed through neural networks with the same dimensionality as the bottleneck layer. Secondly, we used all the genes from the raw data of the pancreas dataset and changed the length of the bottleneck layer.

### 4.4 Experimental details of the single-cell RNA sequencing datasets

#### 4.4.1 Evaluation metric

The baseline metric used in this work is balanced accuracy (bAcc) which is averaged class-wise accuracies. This metric can represent actual class accuracies in a highly imbalanced real-world biological dataset.

Another used metric for evaluating the performance of zero-shot classification is the *h*-score of balanced accuracy for each seen class and unseen class [46].

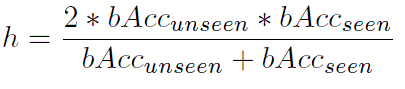

#### 4.4.2 SATL experimental details

In this work we focus on the GZSL task with a transductive setting. The partial data of the target domain is available during training, and the labels are limited to a shared set between the source and target domains. This task was defined and described by a recent survey work by Pourpanah et al. [10]. The data of the source domain and labels are fully utilized during the training. The target data is split into train/test sets, and among the train set, only data of seen classes are available during the training phase. In the test phase, we used a test set with all class labels, including seen and unseen. We evaluate our results with averaged bAcc of all possible combinations of masked labels. For example, the human-mouse pair datasets have eleven classes, thus the average score is calculated on all 55 masking scenarios.

Moreover, prior to applying the HDA method to the sequencing dataset, we need to employ feature extraction methodologies that are widely accepted in the biomedical field. This is equivalent to using pre-trained feature extractors in an image classification task. In particular, we compared Seurat, a typical single-cell sequencing data analysis method with a preprocessing step about highly variable gene selection [35], and DCA [18], unsupervised denoising methods and after highly variable gene selection, with scETM [7], cell topic model and VAE for the feature extraction of single-cell gene expression profiles. Details of the preprocessing and feature extraction are described in the Appendix. With the embedded feature vectors, we searched for the alpha value for CDSPP via class-wise cross-validation [47]. Once we found the best alpha value, we performed the analysis.

#### 4.4.3 Feature importance analysis

The feature importance is obtained by using the explained variance from PCA and the weights of the projection matrix for each species. With this approach, it is possible to investigate which gene contributes more to the common latent space for cross-species analysis in the SATL model. After obtaining the list of genes, we sorted them based on the absolute value of weight and obtained the top 100 contributing genes. The gene lists were further processed using gene ontology analysis for the GO biological process. We used ShinyGO 0.77 [17] and PantherDB 17.0 [25] for gene ontology analysis.

#### 4.4.4 CADA-VAE for single-cell sequencing dataset

Similar to the original work, we implemented VAE with reconstruction loss, cross-modal reconstruction loss, and distribution-aligned loss [19]. In addition to that, another version of CADA-VAE is implemented in this work. CADA-VAE2 implemented two VAEs for each species since they have different numbers of genes. The CADA-VAE2 with two VAEs also has the same loss function with original work, reconstruction loss, cross-modal reconstruction loss, and distribution-aligned loss. Both models are trained with paired data having the same label from each species. The cross-reconstruction loss is calculated as latent vectors from the hidden layer fed into the other VAE.

#### 4.4.5 Domain homogenization using external knowledge

Orthologous genes information can integrate different datasets from different species. It is viable when orthologous gene information is available, and two species share most of the genes. During this gene conversion process, information loss is inevitable. However, this process can homogenize heterogeneous domains of different species and the label-transfer task becomes more feasible. For the comparison, we employed two recent methods that allow transferring cell types across the species barrier, scNym [23] and scAdapt [5]. Both scNym and scAdapt utilize orthologous gene information. They work in an unsupervised way, not relying on any labeled target domain data. Thus, all target cells are masked. The no new identity setting is chosen for scNym to disable the function to find novel cell types. Furthermore, scNym allows for explicitly handling unbalanced source domain cell types. scAdapt is run with a batch and embedding size of 256, using the proposed deep adversarial network option.

#### 4.4.6 Homogeneous domain adaptation

Domain adaptation methods are tested with the python package ‘ADAPT’ [48]. The compared methods are IWC [49], FA, PRED, TrAdaBoost [50], SA [51], CORAL [52], KMM [53], NNW [54], TCA [55], BW [56], TransferTreeClassifier [57], TransferForestClassifier [58]. The same task schemes are applied—the DA model trained on the source domain and the seen classes’ data of the target domain. During testing, seen and unseen classes are predicted by the trained model.

#### 4.4.7 GZSL-QFSL

The baseline QFSL model without the additional Quasi-Fully Supervised Learning loss is considered for this work. This loss is supposed to alleviate the bias towards seen classes by encouraging high probabilities for unseen classes when samples of these classes are evaluated, utilizing unlabeled data during training. The application of QFSL still makes sense since even the baseline model does achieve competitive results [20].

## Declarations

## Acknowledgments

This work contains the results of Master thesis of NPM under the supervision of YP and AH. Figures 1, 5, and 6 were created with BioRender.com

## Code availability

The main analysis code is available in a GitHub repository, https://github.com/iron-lion/HDA4SATL. The code for a Python implementation of CDSPP for GZSL is available in a GitHub repository, https://github.com/iron-lion/cdspp-hda-py.

## Authors’ contributions

The conceptualization of the study was conceived by YP. and AH. Computational experiments were performed by YP and NPM. YP and NPM generated results. YP prepared figures. YP and AH wrote the manuscript and performed review and editing.

## Appendix A SATL results of three single-cell RNA sequencing dataset

**Table A1.** Average unseen accuracy of SATL of each cell type of Bone Marrow dataset and number of samples for each cell type.

| cell type | human#samples | mouse#samples | *Seurat*: | *DCA*: | *scETM*: | scAdapt | scNym |
| --- | --- | --- | --- | --- | --- | --- | --- |
| Myeloblasts | 1179 | 603 | 0.678 | 0.756 | 0.795 | 0.936 | 0.995 |
| Pro-B | 326 | 225 | 0.142 | 0.277 | 0.294 | 0.377 | 0.368 |
| T-NK | 541 | 60 | 0.412 | 0.496 | 0.838 | 0.939 | 0.994 |
| Monoblasts | 358 | 300 | 0.521 | 0.367 | 0.713 | 0.771 | 0.215 |
| Erythroblasts | 3217 | 2845 | 0.533 | 0.676 | 0.961 | 0.953 | 0.968 |
| Pre-B | 623 | 62 | 0.302 | 0.165 | 0.372 | 0.003 | 0.990 |
| Neutrophils | 467 | 267 | 0.331 | 0.784 | 0.978 | 0.998 | 0.996 |
| HSPCs | 307 | 303 | 0.534 | 0.043 | 0.828 | 0.847 | 0.749 |
| Myelocytes | 1431 | 266 | 0.367 | 0.466 | 0.680 | 0.556 | 0.591 |
| Monocytes | 721 | 217 | 0.769 | 0.097 | 0.894 | 0.768 | 0.872 |
| Pericytes | 64 | 112 | 0.108 | 0.460 | 0.631 | 0.922 | 0.844 |

**Table A2.**
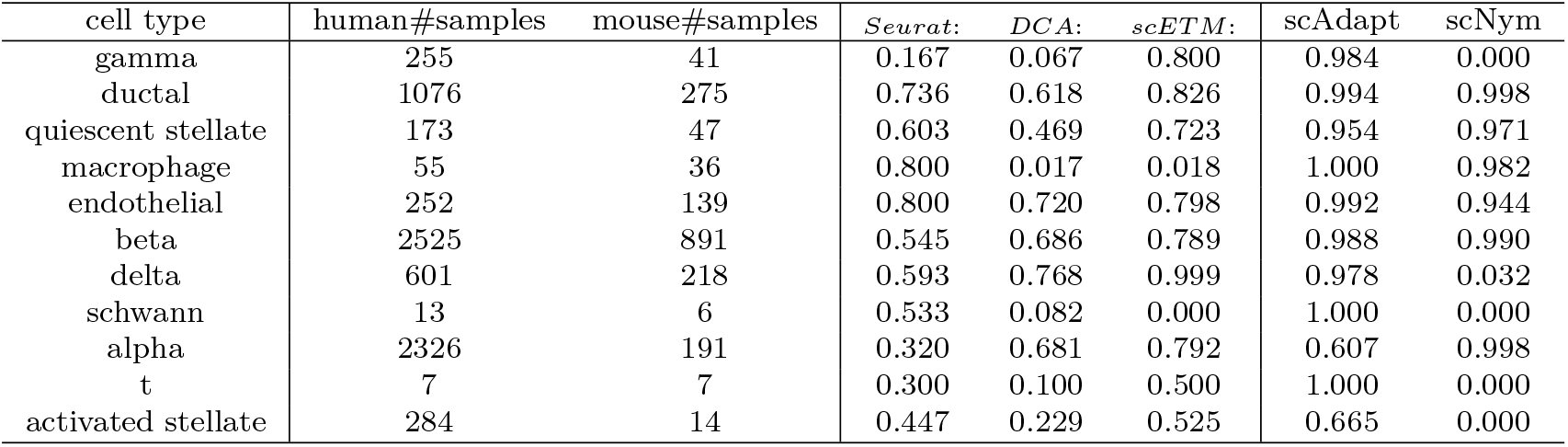
Average unseen accuracy of SATL of each cell type of Pancreas dataset and number of samples for each cell type.

**Table A3.** Average unseen accuracy of SATL of each cell type of Brain dataset and number of samples for each cell type.

| cell type | human#samples | mouse#samples | *Seurat*: | *DCA*: | *scETM*: | scAdapt | scNym |
| --- | --- | --- | --- | --- | --- | --- | --- |
| Lamp5 | 554 | 752 | 0.642 | 0.200 | 0.525 | 0.917 | 0.993 |
| Sncg | 123 | 267 | 0.929 | 0.787 | 0.750 | 0.992 | 0.000 |
| Vip | 633 | 896 | 0.514 | 0.462 | 0.855 | 0.946 | 1.000 |
| Pvalb | 949 | 605 | 0.697 | 0.500 | 0.873 | 0.960 | 0.976 |
| L6b | 245 | 322 | 0.800 | 0.600 | 0.998 | 0.988 | 0.000 |
| Sst | 746 | 850 | 0.701 | 0.571 | 0.900 | 0.992 | 0.976 |
| L6 IT | 228 | 819 | 0.102 | 0.654 | 0.609 | 1.000 | 0.000 |
| L2/3 IT | 3074 | 1526 | 0.770 | 0.599 | 0.791 | 0.780 | 0.981 |
| L5 IT | 1721 | 1320 | 0.756 | 0.783 | 0.739 | 0.911 | 0.940 |
| L5/6 NP | 180 | 510 | 0.497 | 0.807 | 1.000 | 0.989 | 0.983 |
| L6 CT | 455 | 2876 | 0.700 | 0.824 | 1.000 | 1.000 | 0.000 |

## Appendix B Semi-supervised setting comparison

The comparison with other heterogeneous domain adaptation methods is done under a semi-supervised setting. LPJT [21] and DDA [22] were compared with our SATL. During the training, the source data is all used and target data is used only five samples per class.

Detailed parameters of locality preserving joint transfer model (LPJT).

**Table B4.** LPJT parameters for a semi-supervised task with single-cell sequencing dataset.

| Parameters | T | nu | Dim | inter K | intra K | delta | gamma | mu | weightMode |
| --- | --- | --- | --- | --- | --- | --- | --- | --- | --- |
| Value | 5 | 0.5 | 50 | 5 | 5 | 0.5 | 0.5 | 0.01 | Binary |

Detailed parameters of discriminative distribution alignment model (DDA).

**Table B5.** DDASL parameters for a semi-supervised task with single-cell sequencing dataset.

| Parameters | T | beta | tau | lambda | sub Dim |
| --- | --- | --- | --- | --- | --- |
| DDASL | 5 | 0.001 | 0.002 | 0.001 | 100 |
| DDACL | 5 | 0.001 | 0.002 | 0.001 | 100 |

The semi-supervised setting is that all the source data is available during the training, and the target data is available in only five samples per class. SATL performed the best in the pancreas and brain datasets under the semi-supervised setting, with a drop in overall performance when oversampling was excluded. LPJT and DDA (DDASL and DDACL) were tested without the oversampling method.

**Table B6.** Comparison of semi-supervised task accuracy with different feature extraction methods. Five samples per class were available during the training phase.

|  | BM | Pancreas | Brain |
| --- | --- | --- | --- |
| *Seurat*:SATL<sub>(semi)</sub> | 0.853 | **0.989** | 0.945 |
| *DCA*:SATL<sub>(semi)</sub> | 0.882 | 0.847 | **0.947** |
| *scETM*:SATL<sub>(semi)</sub> | 0.872 | 0.964 | 0.946 |
| *Seurat*:SATL<sub>(semi-over)</sub> | 0.727 | 0.775 | 0.918 |
| *DCA*:SATL<sub>(semi-over)</sub> | 0.823 | 0.672 | 0.915 |
| *scETM*:SATL<sub>(semi-over)</sub> | 0.687 | 0.641 | 0.907 |
| *Seurat*:LPJT<sub>(semi)</sub> | 0.600 | 0.745 | 0.945 |
| *DCA*:LPJT<sub>(semi)</sub> | 0.745 | 0.691 | 0.909 |
| *scETM*:LPJT<sub>(semi)</sub> | 0.310 | 0.345 | 0.564 |
| *Seurat*:DDASL<sub>(semi)</sub> | 0.695 | 0.951 | 0.847 |
| *DCA*:DDASL<sub>(semi)</sub> | **0.900** | 0.881 | 0.852 |
| *scETM*:DDASL<sub>(semi)</sub> | 0.790 | 0.873 | 0.877 |
| *Seurat*:DDACL<sub>(semi)</sub> | 0.827 | 0.872 | 0.894 |
| *DCA*:DDACL<sub>(semi)</sub> | 0.841 | 0.808 | 0.896 |
| *scETM*:DDACL<sub>(semi)</sub> | 0.828 | 0.873 | 0.809 |

BM: Bone marrow dataset

## Appendix C SATL feature investigation

### C.0.1 Ablation test of SATL features

#### Ratio of seen-to-unseen classes

**Fig. C1.**
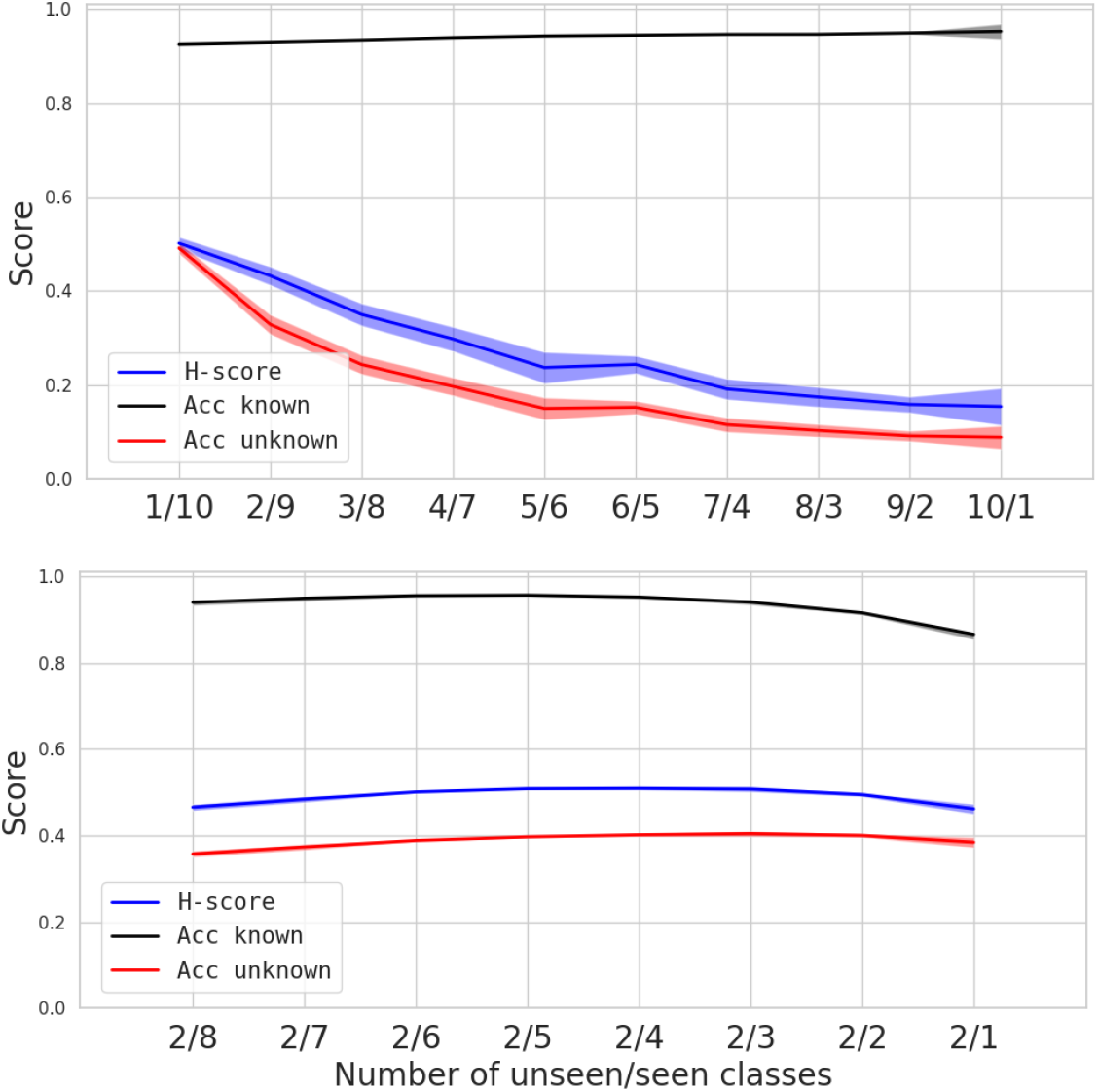
Overall performance varying ratio of seen to unseen classes. The feature is extracted using PCA50. The upper chart shows the total number of classes while varying the ratio of seen and unseen classes. The lower chart shows the number of unseen classes as two and varies the number of seen classes. The overall performance varies with the ratio of seen to unseen classes.

To test the robustness of the SATL method, we investigated how much information is required to achieve zero-shot classification of unseen classes. As the SATL method is trained in a supervised manner with given seen classes, it preserves local structures during projection. We varied the ratio of seen and unseen classes in two different ways, while feature extraction was performed using PCA50. The first test started with a 1:10 split, and the number of seen classes was changed from ten to one. As expected, the task became more difficult as the number of unseen classes increased and the number of seen classes decreased (Figure C1 Upper panel). The second test focused on changing the number of seen classes, while keeping the number of unseen classes fixed at two. Surprisingly, the accuracy of the unseen classes remained fairly stable (Figure C1 Lower panel). These results suggest that the difficulty of the GZSL task with SATL is primarily determined by the number of unseen classes, and that there is no significant benefit to increasing the number of seen classes. Furthermore, we observe that the performance of SATL is more sensitive to the feature extraction method and the number of unseen classes, than to the number of seen classes in the cell-type prediction task.

#### Sampling method

We test the impact of oversampling and pseudo-labeling on SATL (Table C7 in Appendix). When pseudo-labeling is excluded, the overall h-score drops significantly in most scenarios. The pancreas data with Seurat preprocessing cannot show any significant performance improvement. However, this step improves performance in all cases using DCA and scETM. When oversampling is excluded, bone marrow with Seurat and DCA shows a significant h-score drop. In comparison, in the case of pancreas with Seurat or bone marrow with scETM, there is no significant difference. Overall, we observe that this pseudo-labeling and sampling method help improve and stabilize the performance of SATL. However, the effectiveness depends on the dataset and feature extraction methods. There are no detrimental effects of these steps on SATL.

An ablation test was conducted to evaluate the impact of features implemented in SATL. The results show the averaged h-score (refer to Table C7). We tested three single-cell datasets and three feature extraction methods. With DCA and scETM features, sampling with pseudo-labeling demonstrated the best accuracy in all three datasets. However, in the case of the BM and Pancreas dataset with Seurat preprocessing, pseudo-labeling had less impact on performance. In summary, the impact of pseudo-labeling and over-/under-sampling of SATL also depends on the feature extraction method.

**Table C7.** Ablation of SATL analysis comparisons The h-score is shown.

|  | BM | Pancreas | Brain |
| --- | --- | --- | --- |
| *Seurat*:SATL | 0.538 | 0.668 | 0.785 |
| *Seurat*:SATL<sub>(-pl)</sub> | 0.487 | 0.664 | 0.673 |
| *Seurat*:SATL<sub>(-os,-pl)</sub> | 0.328 | 0.607 | 0.631 |
| *DCA*:SATL | 0.535 | 0.541 | 0.710 |
| *DCA*:SATL<sub>(-pl)</sub> | 0.430 | 0.468 | 0.275 |
| *DCA*:SATL<sub>(-os,-pl)</sub> | 0.299 | 0.430 | 0.608 |
| *scETM*:SATL | 0.796 | 0.765 | 0.835 |
| *scETM*:SATL<sub>(-pl)</sub> | 0.533 | 0.389 | 0.632 |
| *scETM*:SATL<sub>(-os,-pl)</sub> | 0.496 | 0.335 | 0.575 |

pl: pseudo-labeling os: oversampling

The impact of the sampling method is evaluated in the same way in a semisupervised task (refer to Table B6). In the semi-supervised task, over-and under-sampling improve accuracy in all three different feature extractors and datasets. Without a sampling method, other HDA methods may show better accuracy.

#### Evaluation of K value

The evaluation of value K was carried out in all nine scenarios and the corresponding h-score changes were visualized in Figure C2. The influence of K was found to vary depending on the feature extraction method employed. In the case of scETM features, the effect of K was minimal, while for Seurat, it showed greater variability but still had only a marginal effect. However, with DCA, the performance improved slightly. These results suggest that the performance of the model is more dependent on the feature extraction method used rather than on the K value. Nevertheless, it is worth noting that increasing K can enhance the model’s performance depending on the feature extraction method utilized.

**Fig. C2.**
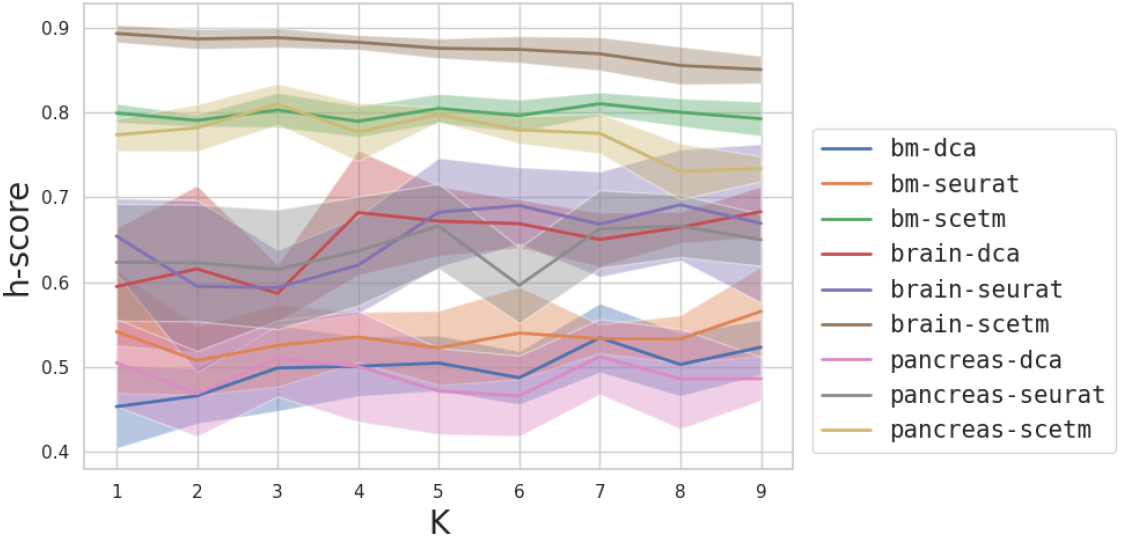
SATL results change according to different K for pseudo-labeling. The shown h-score is obtained from three different datasets and three different preprocessing methods.

